# Developmental models reveal the role of phenotypic plasticity in explaining genetic evolvability

**DOI:** 10.1101/2020.06.29.179226

**Authors:** Miguel Brun-Usan, Alfredo Rago, Christoph Thies, Tobias Uller, Richard A. Watson

## Abstract

Biological evolution exhibits an extraordinary capability to adapt organisms to their environments. The explanation for this often takes for granted that random genetic variation produces at least some beneficial phenotypic variation for natural selection to act on. Such genetic evolvability could itself be a product of evolution, but it is widely acknowledged that the immediate selective gains of evolvability are small on short timescales. So how do biological systems come to exhibit such extraordinary capacity to evolve? One suggestion is that adaptive phenotypic plasticity makes genetic evolution find adaptations faster. However, the need to explain the origin of adaptive plasticity puts genetic evolution back in the driving seat, and genetic evolvability remains unexplained. To better understand the interaction between plasticity and genetic evolvability, we simulate the evolution of phenotypes produced by gene-regulation network-based models of development. First, we show that the phenotypic variation resulting from genetic and environmental change are highly concordant. This is because phenotypic variation, regardless of its cause, occurs within the relatively specific space of possibilities allowed by development. Second, we show that selection for genetic evolvability results in the evolution of adaptive plasticity and *vice versa*. This linkage is essentially symmetric but, unlike genetic evolvability, the selective gains of plasticity are often substantial on short, including within-lifetime, timescales. Accordingly, we show that selection for phenotypic plasticity is, in general, the most efficient and results in the rapid evolution of high genetic evolvability. Thus, without overlooking the fact that adaptive plasticity is itself a product of genetic evolution, this shows how plasticity can influence adaptive evolution and helps explain the genetic evolvability observed in biological systems.

## Introduction

Understanding how evolution works is not complete by understanding natural selection; we also need to understand the generation of the phenotypic variation that natural selection will act on (Hallgrimsson and Hall 2005; Houle et al. 2010). This is challenging since developmental processes often make non-directed genetic variation give rise to phenotypic variation that is highly structured, with some variants appearing more frequently than others (Alberch 1982; Salazar-Ciudad et al. 2003; Kavanagh et al. 2007). If this developmental bias was aligned with the adaptive demands imposed by local environments (i.e., if the mutationally more accessible phenotypes are also the more adaptive ones), adaptive evolution would be greatly facilitated. Such genetic evolvability could itself be a product of past evolution (Uller et al. 2018), but the idea that natural selection would be able to improve genetic evolvability is problematic because the immediate selective gains of evolvability are small on short timescales (Wagner and Altenberg, 1996; Watson 2020).

One suggestion is that the high genetic evolvability is acquired through the capability of organisms to rapidly adapt to their environment during their lifetime (Crispo 2007). However, while adaptive plasticity often appears to ‘take the lead’ in adaptive evolution (e.g., Radersma et al. 2020), the idea that adaptive plasticity explains rapid genetic evolution overlooks the need to explain the origination of the adaptive plasticity that supposedly ‘came first’. In this paper, we seek to better understand whether phenotypic plasticity can help to explain genetic evolvability without overlooking the fact that adaptive plasticity is itself a product of genetic evolution (Levis and Pfennig 2016).

Our approach follows the observation that the phenotypic consequences of genetic variation and environmental variation are not expected to be independent because the consequences of any such variations are channelled by the same underlying developmental mechanisms (Cheverud 1982; Salazar-Ciudad 2006). From this it follows that, if selection for phenotypic plasticity alters the structure of developmental interactions, this will alter how phenotypes respond to genetic mutations and hence affect evolvability. Population genetic models demonstrate that selecting for plasticity can increase genetic variation along the dimensions of the phenotype that are plastic (Ancel and Fontana 2000; Hansen 2006; Draghi and Whitlock 2012; Furusawa and Kaneko 2018). This seems to support a role for plasticity in shaping genetic evolvability. However, given that the interdependence of plasticity and evolvability on development is essentially symmetric, the reverse may be equally possible. That is, if selection for genetic evolvability alters the structure of developmental interactions, this should also alter how phenotypes respond to environmental variation. It thus remains an open question whether genetic evolvability is predominantly shaped by plasticity or *vice versa* (Schwander and Leimar 2011; Levis and Pfennig 2016).

To address this question, we study the relationship between plasticity and evolvability by representing these phenomena in a common framework where the phenotypic effects and adaptive consequences of genetic and environmental variation can be compared. The phenotype distribution that is generated by genetic variation can be represented as a genotype-phenotype (GP) map: an idealized representation of development that assigns a phenotype to each genotype (Alberch 1982; Kirschner and Gerhart 1998; Jimenez et al. 2015). Genetic evolvability is high when the GP map enables genetic change to produce phenotypes that are suitable for adaptation. Analogous to the GP map, plasticity can be understood as an Environment-Phenotype (EP) map (aka reaction norm) that associates each environment with its corresponding phenotype (West-Eberhard 2003; Salazar-Ciudad 2007; Li et al. 2018). Plasticity is adaptive when the EP map enables individuals to produce phenotypes that fit the requirements of the environment in which it finds itself. However, the GP and EP maps do not exhaust the sources of phenotypic variation. Development is also sensitive to non-genetic and non-environmental initial conditions such as biochemical templates, resources and nutrients that are provided by the parents (Newman and Muller 2000; Jablonka and Lamb 2005). The association of such elements with phenotypic variation is often referred to as parental effects (Badyaev and Uller 2009). Here we emphasize the analogous role of such initial conditions in structuring the phenotype distribution by referring to it as the PP (for ‘Parental-Phenotype’) map: a mapping that associates a phenotype with the parentally inherited initial conditions required to produce it.

To model the potential interdependence of these three maps (GP, EP and PP), we use several different and widely used models of development based on gene regulatory networks (GRN). This approach means that we neither assume that the three maps are independent nor that they are related; rather, these are hypotheses we can test. To do this we apply the three different forms of variation (i.e. genetic, environmental, parental) to the same developmental system (GRN), and compare the resulting phenotypic effects. In our approach, genetic changes alter the regulatory connections of the network, environmental changes alter the gene expression levels, and parental effects alter the initial gene expression levels of the network (see Methods). Together, these variables define the GRN dynamics which, when implemented and iterated in the models, result in measurable phenotypes.

In principle, it could be the case that any concordance between the three maps in this model could be intrinsic to the properties of GRNs and arise without selection, or observed only in GRNs subjected to particular selective conditions, or not observed at all. Likewise, it is not known whether selection can, for example, change the GP map produced by a GRN without altering the EP or PP maps, or more generally, whether the effects of selecting for one map has consequences for the properties of the others. Lastly, even if it is the case that selection for one map can determine the evolution of any other map, it could be the case that one of the maps is easier to change with selection than the others. Therefore, the aims of this paper are 1) to model the potential interdependence of the GP, EP and PP maps; 2) to determine how the evolution of one affects the evolution of the other, and whether this evolution is symmetric; and 3) to establish whether or not the three maps are equally responsive to selection.

## Methods

To model how the GP, EP and PP maps can arise and interact, it is necessary to represent developmental systems in a way that allow for genetic, environmental and parental inputs. We therefore expand on three different and widely used GRN-based models of development, each one entailing a different complexity and biological realism (Fig. 1A-C). From the simpler to the more complex, these models are:

**Fig. 1.**
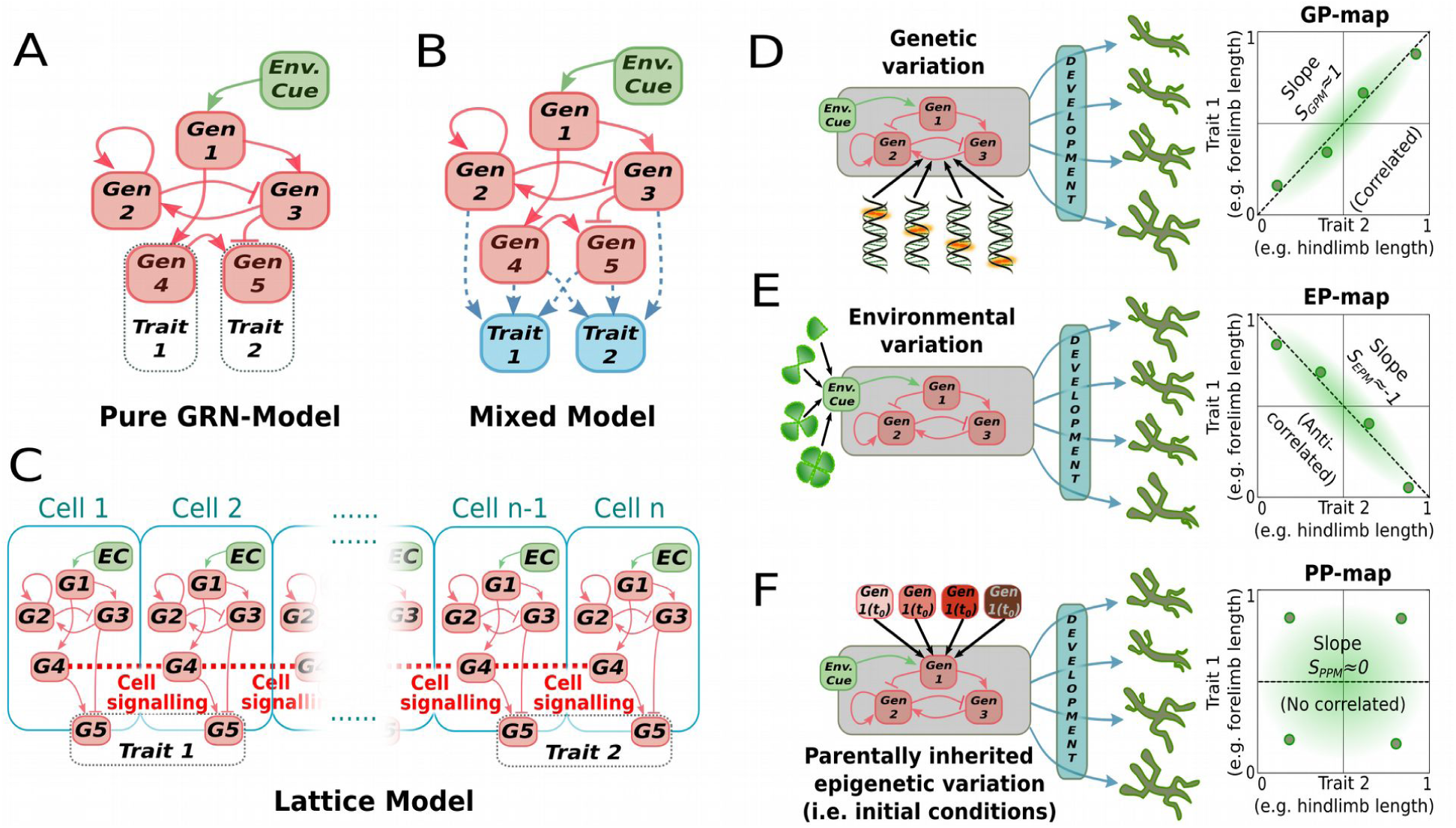
Experimental overview. Conceptual depiction of the three GRN-based models used in this work: **(A)** A pure GRN model where the (two-trait) phenotype is the steady-state concentration of two arbitrary genes. **(B)** GRN + Multilinear model, where each phenotypic trait is calculated as the weighted sum of all the elements within the steady-state GRN. **(C)** Lattice model, where the phenotype is conceptualised as the steady-state expression pattern of one of the constituent genes (Gen 5 in this example) along a one-dimensional row of cells that can communicate between them through cell-cell signalling. In all of these models, phenotypic variation is created by perturbing one or several elements in the core GRN: Perturbations can be introduced in the strength of gene-gene interaction (i.e. as genetic mutations, **D)**; in some environmental cue may regulate some environmentally-sensitive gene **(E)**; or in the (maternally inherited) initial concentrations of each GRN element **(F)**. Perturbations on each of these three different sources of phenotypic variation (one element of the GRN perturbed at a time) will produce a collection of two-trait phenotypes (i.e., hind- and fore-limb lengths). If these phenotypes are plotted in a two-trait (*T*_*1*_*-T*_*2*_) morphospace, they can reveal the structure of the parameter-to-phenotype maps (D-F, right panels). The linear slopes of these maps can be used as a coarse description of these maps, allowing for map-to-map comparisons of random (Fig. 2) and evolved GRNs (Figs 3-5).

### -Basic GRN model

This model (based on Wagner 1994) represents a simple gene regulatory network (GRN; Fig. 1A). It consists of *N*_*g*_ transcription factors that have continuous, positive concentrations (vector *G=(g*_1_,…,*g*_*Ng*_*); gi≥0* ∀ *i*), and regulate the expression of each other by binding to cis-regulatory sequences on gene promoters. The regulatory interactions of this GRN are encoded in the *N*_*g*_ *x N*_*g*_ matrix *B*, whose elements *B*_*ik*_ represent the effect of gene *k* on the transcription of gene *i*. Positive elements (*B*_*ik*_*>0*) represent activation and negative elements (*B*_*ik*_*>0*) represent inhibition. A binary (0 or 1) matrix *M* (*N*_*g*_ *x N*_*g*_) encodes the GRN topology, so that the interaction *B*_*ik*_ is only active if *M*_*ik*_*=1*. The initial state of the vector *G* (represented by the vector *G*_*0*_) accounts for the initial state at the beginning of development, which is supposed to be parentally determined. The vector *E* contains *N*_*e*_ environmental factors (*N*_*e*_*≤N*_*g*_) which affect the levels of gene expression by activating or repressing them (with an intensity of *0≤E*_*i*_*≤1*).

Developmental dynamics are attained by changes in gene concentration over a number of developmental iterations (*t*_*dev*_), and the phenotype is recorded as the steady-state expression levels of two arbitrarily chosen genes in *t*_*dev*_ (Fig. 1A). Only viable (temporally stable) phenotypes are considered: normalized *G, G\**, must remain the same within a threshold of *10*^*-2*^ over an interval of *t*_*dev*_/10 developmental time units (|*G\**_*0*.*9·tdev*_*-G\**_*tdev*_|*≤10*^*-2*^). The gene-gene interactions within the GRN follow a non-linear, saturating Michaelis-Menten dynamics (a special type of Hill function), so that the concentration of the gene *i* changes over developmental time according to the following differential equation:

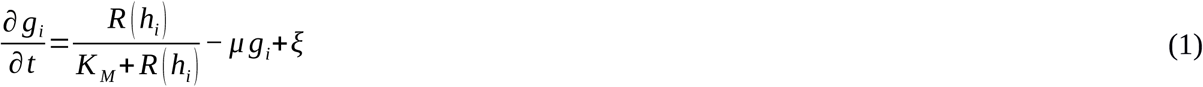

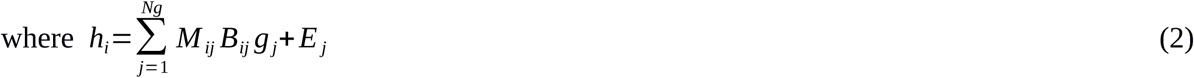

and *R (h*^*i*^) is the Ramp function (*R(x)=x*, ∀ *x≥0* and 0 otherwise) which prevents negative concentrations in gene products resulting from inhibiting interactions, and *K*_*M*_ is the Michaelis-Menten coefficient. Without loss of generality, we set *K*_*M*_=1 (other choices of *K*_*M*_ or specific Hill functions are known not to affect the results, see Salazar-Ciudad et al. 2000). Eq. (1) was numerically integrated using the Euler method *(δ*_*t*_*=10*^*-2*^). All genes and gene products (but not environmental factors) are degraded with a decay term *μ=0*.*1*. Noise is introduced in the system through the term *ξ∼U(−10*^*-2*^, *10*^*-2*^).

### -GRN + Multilinear model

This model (based on Draghi and Whitlock 2012), can be viewed as a multi-linear model of phenotypic determination (Hansen 2006) that is added to a basic GRN-model (Wagner 1994) (Fig. 1B). The key difference with the previous model lies in how each phenotypic trait is generated. Rather than being the expression level of one element of the GRN, each trait *T*_*i*_, *i=(1,2)* receives a contribution from each transcription factor according to a linear coefficient:

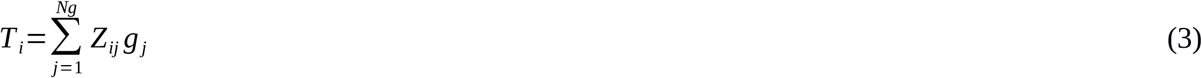

where the factor *Z*_*ij*_ represents the contribution of the *j*^*th*^ gene to the *i*^*th*^ trait (*-1<Z*_*ij*_*<1*). Note that the *Z* matrix encoding the linear coefficients is separated from the matrix *B* encoding the GRN itself. In this paper, the evolutionary implications of the correlations between maps are reported on the basis of this model.

### -Lattice model

This reaction-diffusion model (based on Salazar-Ciudad et al. 2000 and Jimenez et al. 2015) represents a simple developmental model that implements multicellular phenotypes in an explicitly spatial context (Fig. 1C). The model describes on a one-dimensional row of *N*_*c*_ non-motile cells (*N*_*c*_*=*16 in our case), whose developmental dynamics is determined by a GRN (as described in the basic model) that is identical for all cells. Interaction between the different cells is achieved through cell-cell signalling involving extracellular diffusion of morphogens (*N*_*g*_*/3* of the GRN elements are considered to be diffusible morphogens). Each of these morphogens has a specific diffusion rate *D*_*i*_ (*0<D*_*i*_*<1*) and follows Fick’s second law. Zero-flux boundary conditions are used. Thus, the concentration of gene *i* over developmental time now is calculated as:

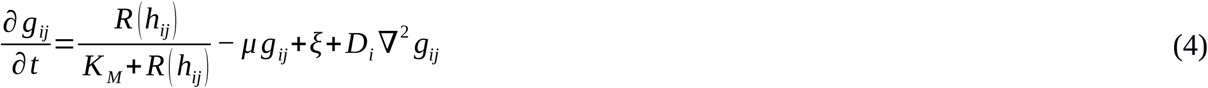

In most works that use this model, the phenotype is conceptualised as the expression pattern of one of the constituent genes along the row of cells (e.g. Salazar-Ciudad et al. 2000; Jimenez et al. 2015). Here, for the sake of comparability with the other models used, we set *T*_*1*_ and *T*_*2*_ as the average concentration of gene 1 in the first and last two cells of the organism (*T*_*1*_*=(g*_*1,1*_*+g*_*1,2)*_*/2*, and *T*_*2*_*=(g*_*1*_,_*Nc- 1*_*+g*_*1*_,_*Nc*_*)/2)*.

While these models differ in complexity, all three feature a GRN at their core, which provides an intuitive way to link each of the constituent elements of the GRN to a different source of phenotypic variation: (i) changes in gene-gene interaction strengths in the GRN can be conceptualised as an effect of genetic variation; (ii) changes in the environmentally sensitive elements in the GRN (sensor nodes and diffusion rates) as an effect of environmental variation; and (iii) changes in the initial concentration of each transcription factor in the GRN as an inherited initial state (Fig. 1D-F). This approach exhausts the ways in which a specific GRN can vary (these variations do not alter the GRN topology (i.e. the *M* matrix), which here is assumed to evolve much slower than the inputs, Salazar-Ciudad et al. 2003).

With the described settings, all the models used in this paper produce a single, 2-trait (2-dimensional) phenotype for each combination of inputs. Thus, a set of phenotypes (i.e. a phenotype distribution) can be generated by introducing variation in those inputs. These phenotype distributions represented in a 2D morphospace are considered maps: those resulting from variation in the genetic inputs are GP maps, whereas those resulting from environmental perturbations or perturbations in the initial conditions are considered as EP and PP maps, respectively. The 2D representation of the whole set of phenotypes that a given developmental mechanism can create from the sum of all perturbations is referred to as a general phenotype distribution, or GPD (i.e., the GP, EP and PP maps are all contained within this GPD, Fig. S1).

## Results

### GP, EP and PP mappings are correlated in randomly generated GRNs

We first explore the inter-dependence between GP, EP, and PP maps in a large (*n>10*^6^) ensemble of randomly generated GRNs. We then separately introduced random variation (10 input values 0<x<1) in the genetic, environmental and parental (i.e., initial conditions) inputs of each of these GRNs, and compared the resulting phenotypic distributions, that is, the resulting GP, EP and PP maps.

We estimated the similarity between these three maps by testing whether or not variation in the genetic, environmental or parental inputs produce similar covariation between the two traits (Fig. 1D-F), using the linear slopes in the phenotypic morphospace as basic descriptors of the different maps. Pairwise comparisons between the slopes caused by variation in genetic, environmental, or parental inputs were all significantly positive (Pearson *r* ≥0.3; Fig. 2A). This demonstrates that GP, EP and PP mappings are not independent in random GRNs. Note that this positive correlation between maps does not imply that the map-specific slopes themselves are positive; only that their slopes, which can be positive or negative, are similar (a comparison between maps that does not consider the direction of the slope is necessary since genetic evolvability is concerned with the sensitivity to random mutations, not their direction; Watson 2020). This interdependence across different mappings is stronger for large and densely connected GRNs (Fig. S2), and is robust to more detailed measures of map-to-map similarity, such as Euclidean distances (Fig. S3). GRNs showing zero or negative correlation between different mappings also exist, but they are less frequent (Fig. 2B).

**Fig. 2.**
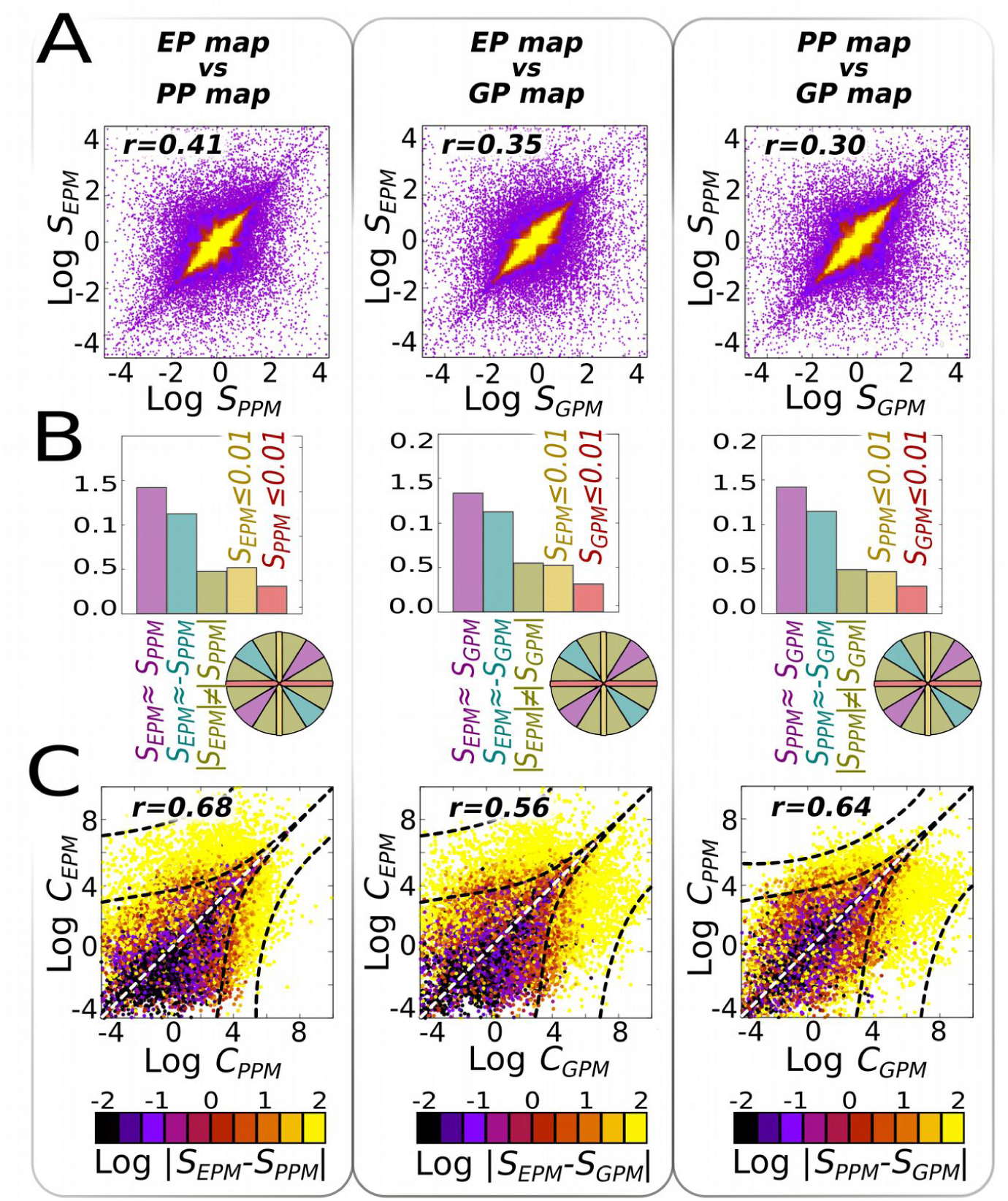
Phenotypic distributions arising from genetic, epigenetic or environmental perturbations are not independent. In a large random ensemble of GRNs (*n=10*^*6*^), systematic parametric variations were introduced into each of their elements. Each perturbation on an element generates a collection of phenotypes in a two-trait morphospace (a *XPM* map), characterised by a linear slope *S*_*XPM*_ (see Fig. 1D-F). **(A)** For each GRN, we compare these slopes, two by two, searching for their correlations in the two-slope spaces (note that these are not two-trait morphospaces). Each dot is a GRN, and the yellow shaded region contains 90% of the GRNs. Correlations are significant (Pearson *r*>0.3) for every combination of maps considered. **(B)** Histograms showing the probability distribution of maps with developmental insensitivity to the first (*Sx*≤0.01; yellow) or second (*Sy*≤0.01; red) type of inputs; and of correlated (parallel slopes); anti-correlated (perpendicular slopes) and non-correlated slopes (otherwise). Each of these cases correspond to the sector of a hypothetical circumference engulfing all points of (A), as exemplified in the coloured circumference, and the relative frequency represents the probability of each point to be located within each sector. **(C)** The complexities of the parameter-to-phenotype maps (i.e., how non-linear they are, see Methods); rather than between their linear slopes are also positive (Pearson *r>0*.*56*). In (C), the colour represents slope similarity: similar slopes (black colour) are associated to simpler (i.e., more linear) maps. *n=30* replicates, GRN + multilinear model (see SI for correlations under other models and Fig. S5 for a null model on C).

To eliminate the possibility that these observed correlations were caused by similarities in the input values, rather than in the structure of the GRNs, we gradually randomize the input parametric values while recording the correlations between maps (Fig. S4). This procedure revealed that the correlations do not depend on particular choices of the input parameters. In contrast, correlations are extremely sensitive to parametric changes in the GRN topology, suggesting that the observed similarity between maps is caused by the structure of the GRN connections (i.e., ‘developmental mechanism’ *sensu* Salazar-Ciudad 2000; Jimenez et al. 2015) rather than the structure of the input perturbations.

### The complexity of GP, EP and PP maps are correlated in randomly generated GRNs

We also investigated whether or not the GP, EP and PP maps exhibit similar complexity in random GRNs. We defined map complexity as the degree of non-linearity in phenotypic response to inputs. This captures the intuition that a linear slope is less complex than a U-shaped response, which is itself simpler than a W-shaped response. Comparing the map complexities between the *10*^*6*^ random GRNs reveals that map complexities are, on average, positively correlated (Pearson *r*≥0.56; Fig. 2C and S2). In other words, if a map (e.g., GP) is simple, the other maps (EP and PP) will be simple too, and they will exhibit very similar slopes. In contrast, if a map is complex, other maps too are likely to be complex, and their slopes will be less similar (Figs. 2C and S2).

To ensure that these observed correlations between map complexities are not a general property of pairs of input-output maps, we analysed a large ensemble of random mathematical functions (polynomials of known degree ≤4) using the same tools that we used for calculating map complexity. This analysis verified that the correlations do not arise between pairs of randomly selected functions unless they belong to the same complexity class (polynomial degree) (Fig. S5).

How a network topology creates similarity between map slopes and complexities can be better understood by looking at the whole set of developmentally attainable phenotypes (general phenotypic distribution: GPD), which can be revealed by means of massive and unspecific parametric perturbations (see Methods and Fig. S1). This procedure shows that each generative network creates a distinctive GPD with a highly anisotropic and discontinuous structure. This structure forces individual maps to be aligned in the same direction simply because many phenotypic directions of change are either very unlikely or developmentally impossible (Figs. S1, S2, S4 and S5).

Positive map-to-map correlations in both slopes and map complexities were found in all three considered models of phenotypic determination (Fig. S2). However, the correlation coefficients are higher and more variable for complex models involving more than pure-GRN dynamics (Figs. 1B-C and Fig. S2).

### Evolving only one of the GP, EP or PP maps changes the phenotypic biases across the other maps

After exploring the map-to-map correlations in random GRNs, we next wanted to address whether or not adaptive changes within one map (i.e., changes in the covariation between traits) are able to induce similar changes to the other maps. To do so, we performed three sets of selection simulations using the developmental model of intermediate complexity (GRN + multilinear). In each simulation, we allowed only one of the three different maps (henceforth the “selected map”) to evolve in response to selection. We refer to the other maps as the “non-selected” maps (see SI for details).

At each evolutionary time step, we introduce variation only in the input associated with the selected map (i.e., genetic, environmental or parental inputs). In response to that variation, each individual develops a set of phenotypes that is compared to an arbitrary (linear) target map to determine the individual’s fitness. Thus, the entire phenotype distribution produced by the selected map is accessible to natural selection (i.e., fine-grained selection). On the contrary, the inputs of the non-selected maps were kept fixed (no variation) during simulations, so that these maps remain effectively “invisible” to natural selection (Fig. 3A). Once each evolutionary simulation reached a steady state, we assessed if there were any changes in the non-selected maps. We did this by collapsing the variation for the selected map to a single input value (*x∼U(0,1)*) and introducing variation to each of the non-selected maps. This experimental setup guarantees that any observed changes in non-selected maps can be attributed to indirect effects of direct selection on the selected map.

**Fig. 3.**
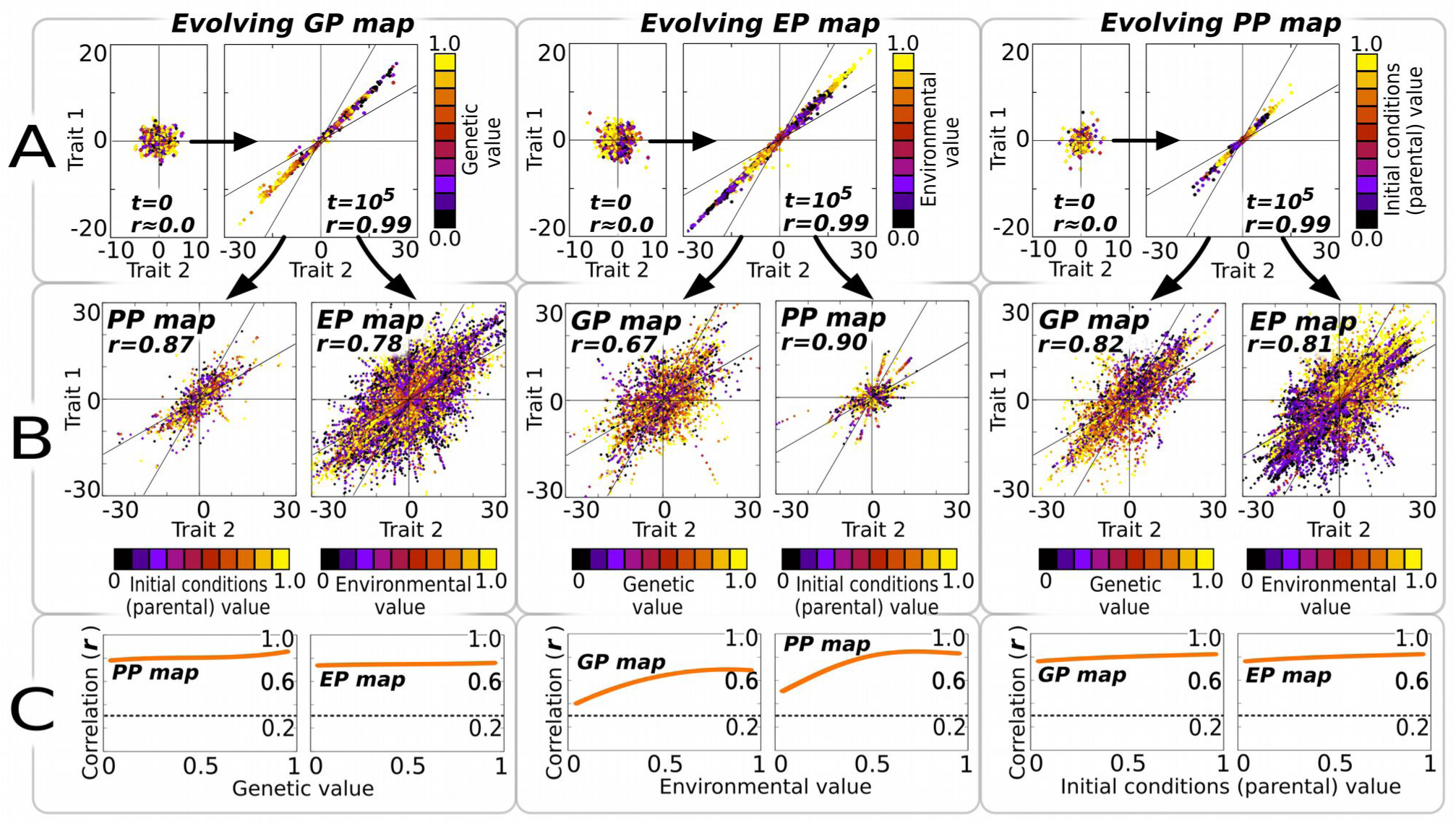
Evolving a single map creates similar phenotypic distributions in the other maps. A population whose individuals initially exhibit a random phenotypic distribution in *t=0* **(A**, small panels) is evolved to fit a target phenotypic distribution (*S*^*T*^*=1*) using as an input just one kind of phenotypic determinant (i.e., genetic, environmental, or parental variation). Other targets (*S*^*T*^*=−1*) give similar results (see Fig. S6). In each generation, one individual is exposed to 10 different input values (*0<x<1*) of a single phenotypic determinant (the colour of each dot in A-B represents value of this input). This parametric variation produces a set of ten potential phenotypes whose slope is compared to the target to evaluate the individual’s fitness (see Methods). After *10*^*5*^ generations in a mutation-selection-drift scenario (where other sources of phenotypic variation are frozen), the population has a narrow phenotypic distribution in the evolved map (A, large panels). In **(B)** we uncover variation in the other maps by introducing parametric variation (*0<x<1*) in the phenotypic determinants that were kept fixed during the evolutionary trial. Results reveal that selection on a single map creates significant side-effect phenotypic distributions in the other maps that are not the target of selection. **(C)** Correlations in the side-effect maps are significant across all parameter values at which the parameter of the evolved map is frozen. *p=64* individuals; *n=30* replicates, GRN + Multilinear model.

The results revealed that evolving any one map modifies the other maps as well, introducing in them the same adaptive phenotypic biases as observed in the selected map (Pearson *r>0*.*3*, Fig. 3A). This holds true for every map combination (Fig. 3B) and across the entire range of parameters we tested (Fig. S2). However, the phenotype biases in the non-selected maps are not as strong (| *r*| *<0*.*5* in some cases) as in the map under selection (*r=0*.*99*), and exhibit substantial temporal variation (Fig. 4). The results in Figure 3 illustrate the outcome of selecting for a linear map with a slope *S=1* in the two-trait morphospace, but simulations with *S=−1* or with changing selective pressures yielded similar results (Figs. 4 and S6).

**Fig. 4.**
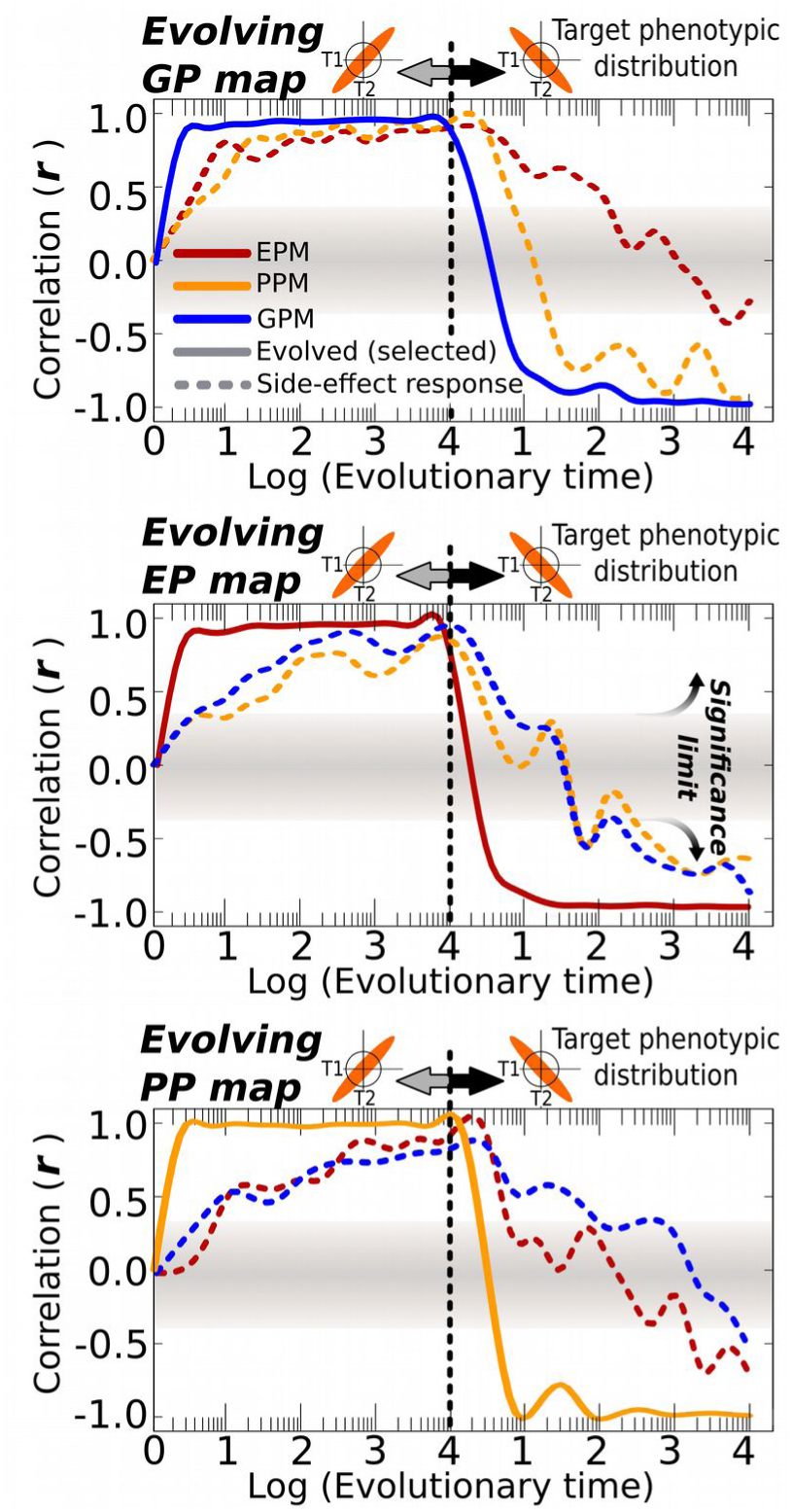
Side-effect phenotypic distributions are able to track shifting targets. These plots show how initially unstructured populations are able to adaptively evolve a specific target distribution (a single map with a defined slope of *S*^*T*^*=1*, solid lines), which creates as a side-effect correlated phenotype distributions in the other maps (dashed lines). The middle point corresponds to the steady-state situation shown in Fig. 3. For these plots, the target slope has been shifted to *S*^*T*^*=−1* at generation *t≈10*^*4*^, showing how the maps that evolve as a side-effect are able to “follow” the one that is being selected. This pattern is similar for every map considered as long as the selective grain is the same. *p=64* individuals; *n=30* replicates, GRN + Multilinear model.

### Maps evolve faster under fine-grained selection than under coarse-grained selection

In the previous experiments, each evolving population was allowed to sample a wide range of genetic, parental or environmental inputs in each generation, and selection therefore acted on a wide range of phenotypic outputs. In other words, we assumed a very fine-grained selection. Several studies show that adaptive plasticity readily evolves when selection is fine-grained (Levins 1966; Via and Lande 1985; De Jong 1995), although it is not essential (Rago et al. 2019). Whether or not a similar effect occurs for GP and PP mappings is unknown. To address this, we explored the ability of every map to adapt to a target map under different levels of selective grain, ranging from very fine-grained selection (where individuals can experience several inputs within their lifetime) to the very coarse-grained cases in which there is just one input per generation and this input only shifts every *n* generations.

As Fig. 5 shows, all maps evolve more efficiently under fine-grained selection than under coarse-grained selection. Furthermore, the ability to adapt to the target map escalates sharply around a grain value of 1 (Figs. 5 and S8). Under the metrics adopted here (see SI), this is the value where single individuals experience on average more than one input per generation. This implies that it is much easier to evolve a map efficiently, and thus to affect the other maps, if there is within-lifetime variation in the inputs to that map.

**Fig. 5.**
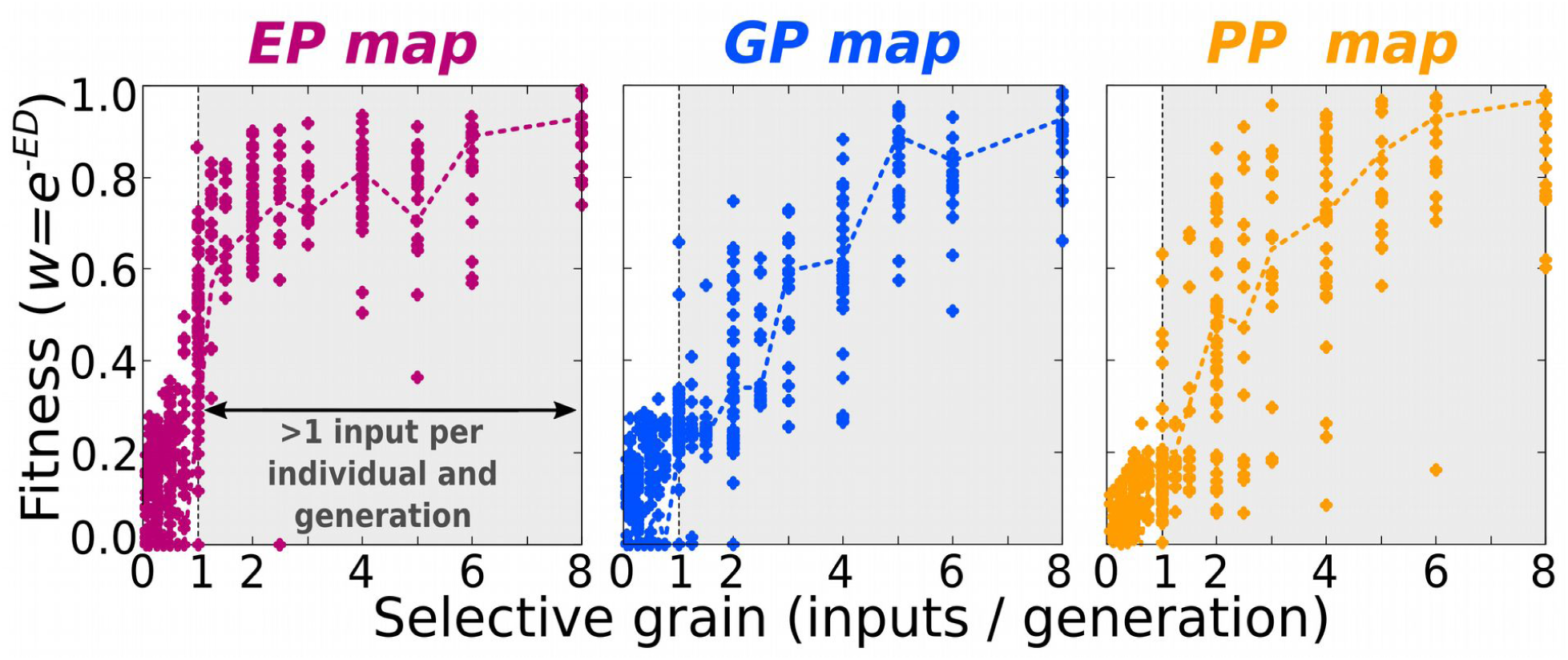
Map evolvability depends on selective grain, and it is maximal for Environment-Phenotype maps. In figures 3 and 4, simulations assumed that natural selection could act on the entire map. This corresponds to very fine-grained selection since we define selective grain as the average number of parameter-phenotype points that be experienced by a single individual in each generation (and hence “seen” by natural selection, see Methods). In this experiment, the assumption about high fine-grainedness has been relaxed. For each level of selective grain, the ability of natural selection to evolve a linear map with an arbitrary slope is recorded as the long-term fitness. Results for environment-phenotype (*EP*) map are depicted in red; for genotype-phenotype (*GP*) map in blue and for parental-phenotype (*PP*) map in yellow. These plots show that the ability to adapt to a target slope increases non-linearly with selective grain, and that maximal efficiency is achieved when selection is fine-grained (>1, shadowed areas), which corresponds to scenarios in which single individuals can experience more than one input per generation. Such high levels of selective grain are typically only attainable for Environment-Phenotype (*EP*) maps (see main text). Points correspond to individual replicates, and dashed lines to averages over the *n=30* replicates. For each replicate, the target map is a linear function of arbitrary non-zero slope. Euclidean-distance (*ED*)-based fitness. *p=64* individuals; *t=10*^*4*^ generations, GRN + Multilinear model.

In most organisms, individuals can experience different environmental inputs during its lifetime but are limited to a single genotype and a single set of parentally inherited initial conditions. As a result, the EP map would be the one most intensely sculpted by natural selection. Because of this, the EP map can exercise a stronger influence on the other maps than *vice versa* when all the three maps evolve simultaneously (Fig. S7).

Besides selective grain, the ability of GRNs to evolve a target map also depends on the complexity of the target map (i.e., how non-linear is the phenotypic response to input variation). Simple (i.e., linear) maps can easily evolve with moderate fine-grained selection (≈2 inputs per lifetime) whilst evolving more complex (i.e., quadratic or cubic) map requires an increasing number of inputs per lifetime (Fig. S8).

## Discussion

Understanding how the processes that generate phenotypic variation interacts with natural selection is necessary to explain and predict the course of evolution (Alberch 1982; Kavanagh et al. 2007; Salazar-Ciudad and Marin-Riera 2013; Uller et al. 2018). While it is easy to understand that any developmental bias aligned with adaptive demands would facilitate adaptation, it is not obvious how these biases originate, nor how they might change or be maintained over evolutionary time. Phenotypic adaptation can precede genetic adaptation, and it has been suggested that plasticity therefore facilitates genetic evolution (reviewed in West-Eberhard 2003). However, trying to explain genetic evolvability by presupposing the existence of adaptive plasticity overlooks the fact that adaptive plasticity is itself a product of genetic evolution. If explaining adaptive plasticity requires past genetic evolution to have already produced adaptive phenotypic responses to particular environmental cues, this does not help to explain genetic evolvability itself. The idea that plasticity and evolvability are intrinsically linked through development provides a way that selection for one can result in the evolution of the other, as studied here. Our aims have been to explore this linkage using mechanistic models of developmental dynamics and thus explore the evolutionary consequences of the relationship between plasticity and evolvability.

Our results show that a concordance between the GP, EP and PP maps is intrinsic to developmental dynamics based on GRNs (with or without selection). Because of this concordance, selection for any map will affect the other maps in an essentially symmetric fashion. However, the efficiency of natural selection in sculpting the different maps is not symmetric: while the immediate selective gains of evolvability are small on short timescales (e.g., to individuals), the selective gains of plasticity can be large. Accordingly, selection for plasticity is much more effective in changing the GRN and genetic evolvability than direct selection for genetic evolvability. Thus, without overlooking the fact that adaptive plasticity is itself a product of genetic evolution, we show how the genotype-phenotype (GP) map can be adaptively shaped by selection for phenotypic plasticity, suggesting that adaptation to environmental variation helps explain the remarkable genetic evolvability of organisms in nature.

The results show that the phenotypic effects of genetic, parental and environmental sources of variation are typically similar across a wide range of assumptions. This finding generalises previous research that found similar correlations on other theoretical grounds or for more restricted scenarios, such as selection for developmental robustness to environmental perturbation (Ancel and Fontana, 2000; Salazar-Ciudad 2006; Salazar-Ciudad 2007; Draghi and Whitlock 2012; Furusawa and Kaneko 2018). Moreover, since the concordance appears in randomly generated regulation networks, it does not require the concourse of past selection to produce this concordance. Rather, stability analysis revealed that correlations between GP, EP, and PP maps are caused by the structure of the network itself and its associated dynamical properties. Although this result might be expected for statistical models that assume linearity and additivity (e.g., the equality *P=GxE* from quantitative genetics, Lande 1982), it is non-trivial for mechanistic models like the ones presented here since these involve non-linear interactions between the inputs. These interactions create a non-uniform space of phenotypic possibilities (a generalised phenotype distribution; GPD) that includes regions of the morphospace that most of the parameter combinations map onto (i.e., phenotypic attractors; Furusawa and Kaneko 2018), and ‘forbidden’ regions that cannot be attained by any parameter combination. These features of the GPD impose strong limitations on the phenotypic variation that is possible, and the shape of the genotype-, environment-, and parental-phenotype maps will be similar since they share the same attractors.

These shared attractors explain why a population that has evolved a specific (e.g., EP) map will show similar biases in all its maps, even when those have not been selected for. However, this dependence would not make plasticity exercise a disproportionate effect on genetic evolvability unless there was some asymmetry that makes selection for properties of the EP map more efficient than selection for properties of the GP or PP maps. We show that this crucial asymmetry follows from differences in the temporal timescale (‘grain’) of environmental, parental and genetic variation that is input to these maps. Specifically, the variational properties of a map evolve faster when individuals experience multiple inputs, and hence can develop multiple selectable phenotypes, during their lifetime (Levins 1966; Via and Lande 1985; De Jong 1995). While individuals can experience multiple environments during their lifetime, they do not experience multiple genotypes or initial conditions (e.g., the distribution of phenotypes produced under genetic variation is a property of a family, population or lineage, not an individual). As a result, selection for GP and PP maps should typically be more coarse-grained and less efficient than for EP maps. This general property of natural selection causes adaptive EP maps to generally evolve more readily than adaptive GP (and PP) maps, even though they depend on the same developmental dynamics.

These results generate predictions can be tested empirically, for example, by means of experimental evolution. One particularly useful approach would be to select populations in environments of different variability (i.e., selective grain), which should result in populations with different EP maps. The prediction is that the finer the selective grain, the more the structure of the GP map will resemble that of the EP map, which can be tested using mutation accumulation experiments or genetic engineering. Whether or not such changes in the GP map changes the capacity for future adaptation could be tested by exposing populations to new selective regimes that are more or less structurally similar to what the population initially adapted to. Other experimental and comparative approaches could also test one or several of the predictions of the relationship between EP, PP, and GP maps (e.g., Lind et al. 2015; Noble et al. 2019; Radersma et al. 2020).

While our results demonstrate that natural selection on phenotypic plasticity would cause the GP map to evolve, they also show that the ability to evolve a certain map is severely limited by the map complexity itself, with complex (e.g., cubic) maps requiring highly fine-grained selection. This would render very complex EP (and GP) maps unreachable by adaptive evolution even in the most fine-grained scenarios. However, complex maps are known to exist, which suggests that other non-selective processes, such as developmental system drift (True and Haag 2001; Jimenez et al. 2015), may play an important role in developmental evolution (Alberch 1982; Salazar-Ciudad and Marin-Riera 2013). This possibility is compatible with our results, which simply state that whenever adaptive developmental biases do exist, they will be predominantly generated through selection for phenotypic plasticity. Furthermore, since each trait needs to maintain its function as other parts of the organism develop and grow, selection for plasticity can be even more fine-grained that expected (our models do not fully capture this developmental dependence because they record the individual fitness after a fixed developmental time).

These results shed light on whether or not plasticity can exercise a predominant role in adaptive evolution, a hypothesis with a long and contentious history in evolutionary biology (see Weber and Depew 2003; West-Eberhard 2003; Schwander and Leimar 2011; Laland et al. 2014; Uller et al. 2019; Parsons et al. 2020). While adaptive modification of environmentally induced phenotypes can make plasticity appear to ‘take the lead’ in evolution without any link between plasticity and genetic evolvability (Radersma et al. 2020), the evolutionary change in the GP map caused by adaptive plasticity suggests that evolution is particularly likely to proceed where plasticity leads. Over longer timescales, this process provides a biologically plausible mechanism for the internalisation of environmental information, resulting in developmental biases whose structure ‘mirrors’ the structure of the selective environment (Riedl 1978), and thereby facilitating further adaptations through genetic modification of environmentally induced phenotypes (Draghi and Whitlock 2012; Watson et al. 2014; Rago et al. 2019; Parsons et al. 2020). Without denying the importance of other (adaptive and non-adaptive) processes, this constitutes a strong argument for a role of plasticity in shaping the path of evolution.

## Acknowledgements

The authors thank J.F. Nash, D. Prosser, J. Caldwell, L.E. Mears, I. Hernando-Herraez and R. Zimm for helpful discussion and comments.

## Author contributions

M.B.U., T.U. and R.A.W. conceived the idea, which was later refined by all the authors. M.B.U., A.R., Ch.T. and R.A.W. established the experimental design. M.B.U. wrote the software and conducted the *in silico* experiments. All authors contributed to the interpretation of the results. M.B.U., A.R., T.U. and R.A.W. wrote the manuscript.

