## Supplementary Information for "Developmental models reveal the role of phenotypic plasticity in explaining genetic evolvability"

1- Institute for life sciences / Electronics and computer sciences. University of Southampton (UK)

2-Department of Biology, Lund University, 22362, Sweden.

*- Corresponding author:

**Correlations between maps.** A large ensemble ($n=10^6$) of random GRNs was created by setting $N_g \sim U(1,24)$ and $p(B_{ij} \neq 0) \sim U(0,1)$, so that a variation in GRN size and connectivity were represented. The same ensemble was used for the three models. Model-specific elements were randomly drawn: $Z_{ij} \sim U(-1,1)$; $D_i \sim U(0,1)$. For each random GRN, parameter-to-phenotype maps were generated through systematic (parametric) perturbations in each of the GRN elements. The element perturbed was randomly chosen for each replicate ($n=30$) and given values from 0 to 1 at 0.1 intervals (in this work, the input values of the different maps are of similar magnitude, an idealisation that allows us to compare the evolutionary properties of the different maps). During these perturbations, GRN topology was always held fixed. Perturbations in $B_{ij}$ were conceptualised as genetic changes; in $G_0$ as changes in the initial conditions (i.e., parental effects); and in $E_j$ or $D_i$ as environmental changes (Fig. 1). That way, the systematic perturbation of each element generated 10 different phenotypes that were recorded in a two trait morphospace, constituting a map (GP map, PP map or EP map, respectively). Note that our “maps” are not maps in a formal mathematical sense because they do not retain the univocal relationship between the inputs and the outputs. However, they allow us to compare different phenotypic distributions whose inputs have different units and magnitudes.

We focus on two-trait phenotypes because they embody the minimal multivariate system where associations between traits can be found. These maps were compared, two by two, using two measures of map-to-map similarity. The first is a coarse-grained measure: Pearson’s *r* correlation between the two linear slopes in the phenotypic morphospace (Fig. 2). In order to take into account negative and close-to-zero slopes, the original slope values were transformed to $S_a=sgn(S)\cdot Log(1+S)$, so that negative values correspond to negative slopes, and not to $0<S<1$ (therefore, the radially symmetric distribution of points around the origin (0,0) observed in in Fig. 2A suggests that individual trait-trait correlations across maps have similar likelihood of being positive or negative). Two maps *a* and *b* were said to be correlated or uncorrelated depending on their sectorial position in this correlational (*Sa*,*Sb*) space: $corr(a,b) \leftrightarrow |tan\text{-}1(Sa/Sb)\text{-}\pi/4| \leq \pi/12$, *anticorr(a,b)* ↔ *|tan-1(Sa/Sb)+π/4|≤π/12,* and not correlated otherwise (Fig. 2B). The second, fine-grained measure is the Euclidean distance ($ED_{a,b}$) between maps *a* and *b* (Fig. S3):

$$ED_{a,b}=\sqrt{\sum_{j=1}^{10}\left(T_{aj1}-T_{bj1}\right)^2+\left(T_{aj2}-T_{bj2}\right)^2} \quad (5)$$

where $T_{ijk}$ is the value of trait *k* in the $j^{th}$ phenotype of map *i*. As Fig. S3A shows, $ED_{a,b}\ \alpha |S_a/S_b|$. As a proxy for map complexity ($C_a$) we use the sum of the squared residuals of each map with respect to its linear regression: the more a map departs from a perfect line the more complex it is:

$$C_a=\sum_{i=1}^{10}\left(T_{i2}-T_{i1}S_a\right)^2 \quad (6)$$

where $T_{ij}$ is the value of the trait *j* of the *i*th point (phenotype) of the map considered. For this analysis maps were re-scaled to ($0<T_{ij}<1$) values in order to avoid size-effects on the map complexity (otherwise the squared residuals of maps with large phenotypic values would result in artefactually higher complexity) (Fig. S3B). We assess the effect on map-map similarity (slopes and map complexities) of GRN size ($N_g$) and connectivity $p(B_{ij}\neq 0)$ (Fig. S4), but not of GRN topology itself as this is beyond the scope of this work (for a discussion on this see Salazar-Ciudad et al. 2000 and Jimenez et al. 2015).

**Control experiments.** Two control experiments were set up to better understand the causes of the observed correlations between slopes *S* and map complexities *C*. In the first, with a probability *p={0.1,0.2,...1}*, GRN topology was changed as $M_{ij}\rightarrow|M_{ij}-1|$ and the GRN input values as $x\rightarrow x\sim U(0,1)$. Then, correlations were recorded between the maps arising from the unperturbed GRN (*S* and *C*) and the randomized ones (*S** and *C**, Fig. S4). In the second control experiment, we used a set of randomly generated mathematical functions (polynomials *f(x)* with known degree *deg(f)≤4*) as a null, non-generative space in which we could test whether the observed correlations between map complexities are a general property of any mathematical function rather than a biologically relevant phenomenon (argument and value of *f(x)* are considered to correspond to the traits $T_1$ and $T_2$). A discrete "mapping" was created by assigning ten values to *x={0.1,0.2,...1}*, and then calculating the corresponding *y*-values (Fig. S5):

$$y\left(\approx T_2\right)=\sum_{i=0}^{4} R_{(i)}\, x\left(\approx T_1\right)^i e^{-i} \quad (7)$$

where $R$ is a vector of random numbers $R(i) \sim U(0,1)$ and $e^{-i}$ a corrective token that devalues the high-degree terms of the function, ensuring that polynomials of different degrees are equally represented. If necessary, $y$-values were rescaled to ($0<y<1$, as in Fig. S3B), so that the map complexity of the the function was measured under the same conditions as for GRNs.

**Evolving maps.** Several of our experiments involve the adaptive evolution of a map: a population of $p(=64)$ haploid individuals picked from the random ensemble evolves in a mutation-selection-drift scenario until a map with a target slope $S^T$ is encountered, or until a maximum number of $t_{max}=10^5$ generations is reached. Arbitrarily, $S^T$ is set to $S^T=1$ (other choices do not alter the results, see Fig. S6). With a rate of 0.05 ($=1/N_{gmax}$) per element and generation, point mutations are introduced in the matrices encoding the topology and interaction strengths of the GRN: $B_{ij} \rightarrow B_{ij}+\xi$ ($\xi \sim N(\mu,\sigma)$; $\mu=0$, $\sigma=0.01$) and $M_{ij} \rightarrow |M_{ij}-1|$. The fitness of each individual $W_i$ is calculated on the basis of its ability to create a map similar to the target one (not on the basis of a single phenotype). Thus, each individual in each generation is exposed to 10 different inputs in one of its GRN elements (the element depends on the map being evolved), and its slope $S_i$ in a $T_1$-$T_2$ morphospace recorded and compared to the target slope $S^T$. This algorithm is formally equivalent to an inter-generational variation in the inputs (De Jong 1995). The similarity with the target slope determines the individual's fitness and, in turn, the probability of each individual to contribute to the next generation:

$$W_i = e^{-|S_i - S^T|} \quad (8)$$

Some of our experiments involve different levels of selective grain on the maps, which has two different components: intra-generational (i.e., how many different inputs (or points of the whole map) can the population experience in a single generation) and inter-generational (i.e., how often these inputs change, which can be conveniently expressed as the number of generations between changes in the input values). For the sake of simplicity we collapse these two components in a single composite measure of fine-grainedness as inputs/generation (Fig. 5, Fig. S7 and Fig. S8). Since slopes alone cannot account for the number of points in a map, the fitness is now calculated as:

$$W_i = e^{-ED_{mapi,map^T}} \quad (9)$$

where $ED_{mapi,mapT}$ is the Euclidean distance, point by point, between the individual's map ($map_i$) and the target map ($map^T$), as described in Eq. (5).

### Supplementary Figures:

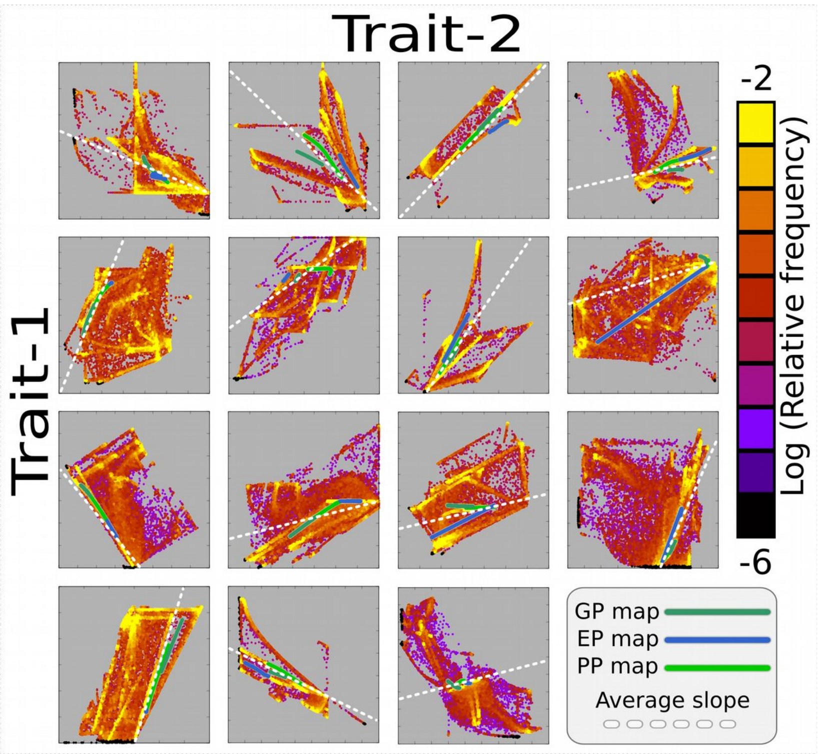


**Figure S1. General Phenotypic Distributions (GPDs).** A general phenotypic distribution (GPD) is the total amount of phenotypic variation that a given developmental mechanism (with a fixed GRN topology) can produce under systematic variations of all their phenotypic determinants (i.e. genetic, parental and environmental perturbations). The concept of GPD explicitly expands the previously proposed concept of "variational properties" (See Ref. 24) in order to incorporate parental variation. This figure shows 15 different GPDs, each one corresponding to a different but representative GRN of our random ensemble. For each one, instead of varying one single GRN element at a time using ten pre-established values (0 to 1 at 0.1 intervals), we vary all of them simultaneously and in a more continuous manner (100 random values $0<x<1$). That is, instead of $n=30$ phenotypes (10 genetic+10 parental+10 environmental perturbations), we get $n=10^6$ (100x100x100) potential perturbations (and phenotypes) per GRN. Each point in these panels corresponds to one of these $10^6$ potential phenotypes in a $T_1$-$T_2$ morphospace (scaled to fit the the minimum and maximum $T_1$-$T_2$ values of the GPD). The colour represents the density of points: regions with high density (calculated on the basis of a 100x100 grid) are coloured in yellow, and regions with rare phenotypes are coloured in purple. Notice that the GPD has two levels of structure: One is the set of developmentally possible phenotypes (the occupied area of the morphospace), which as the panels show, is often discontinuous and non-isotropic. The other is the different likelihood of the different phenotypes within this region (specially the most probable phenotypes exemplified by the yellowish regions and ridges). This two-level structure of the GPD suggests that development creates correlations between the maps (green and blue lines) because the maps most likely contain phenotypes belonging to the ridges of maximal phenotypic probability (which are often parallel or sub-parallel). Dashed white lines are the average slope of the GP, EP and PP maps (dark green blue and light green lines, respectively). GRN + Multilinear model.

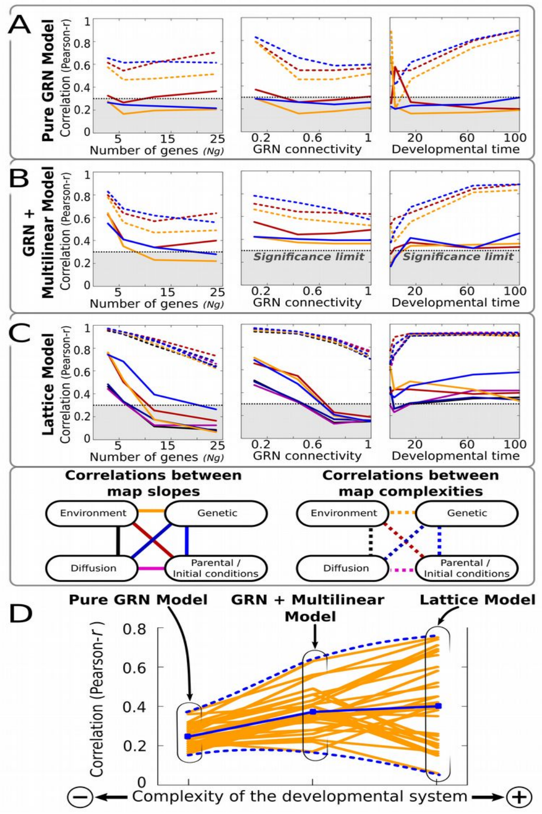


**Figure S2. Map-to-map correlation are robust across the different models considered.** A large ensemble (*$n=10^6$*) of random developmental mechanisms was generated by creating a random GRN (as in Fig. 2, see SI1) and by assigning random values (*U~(0,1)*) to the remaining model-specific parameters (see Methods). For each of these mechanisms, systematic parametric variations were introduced to each of the GRN elements (genetic, initial conditions, environmental inputs). In the case of the lattice model, variation was also introduced in the diffusion rates *D*. Each perturbation on an element generates a collection of phenotypes (a map) in a two-trait morphospace, characterised by a linear slope *S* and a map complexity *C*. The panels show the correlations (Pearson's *r*) between pairs of slopes (solid lines) and pairs of map complexities (dashed lines) for the three models considered: the pure GRN Model (A), the GRN + Multilinear model (B) and the lattice model (C). The schemas below panel (C) assigns a specific colour to each map-map pair (notice that the three models are readily comparable because they map onto the same two-trait morphospace and have similar ranges in the parametric variation). For each model, the randomly generated mechanisms have been sorted according to their number of genes ($N_g$) and GRN connectivity (proportion of non-zero elements in the matrix *B*), to see how these topological features affect the map-to-map correlations (left and central panels). In addition, for each mechanism the correlations have been recorded over a different number of developmental iterations (right panels). Overall, this figure shows that Pearson correlations are significant (*$r>0.3$*) across the different models considered and under different topological and developmental constraints. (D) Since the different mechanisms bear the same (*$n=10^6$*) GRNs, we assess the effect of an increased model complexity in the correlations. Yellow lines show how adding extra layers of complexity (e.g., a multi-linear layer or multicell reaction-diffusion processes) to a basic GRN affects the map-to-map correlations.

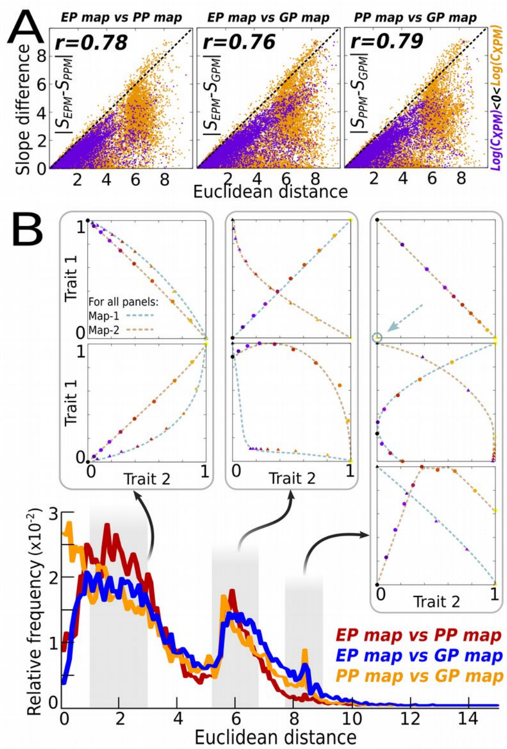


**Figure S3. Correlation between maps is robust when using fine-grained measures of phenotypic distances.** In addition to slope correlation (a coarse-grained measure of phenotypic distances) between maps, we have also measured map-to-map similarity using the euclidean distances (*ED*) between them (a fine-grained measure of phenotypic distances that takes into account distances between pairs of individual phenotypes). (A) *ED* between paired maps for each GRN in our random ensemble are compared with the slope similarity (absolute value of slope differences) between maps. We proceed that way because there is not an *a priori* null expectation about which would be the minimum *ED* value above which the similarity would be significant. The three panels show that slope similarity strongly correlates (Pearson's $r>>0.3$) with the Euclidean distance between any pair of maps. This implies that the former coarse-grained measure can be confidently used as a proxy of map similarity, as it has been extensively used in most plots. In addition, we have detected that complex parameter-to-phenotype maps ($Log(C)>0$), yellowish dots) have a much noisier relationship between slope differences and euclidean phenotypic distances (*EDs*) than simpler maps ($Log(C)<0$), purple dots). This is probably due to the fact that, although complex maps tend to have larger *EDs* between them, their linear slopes tend to be small, which explains the great scattering of yellow dots below the diagonal. (B) The whole random ensemble is analysed now in terms of *EDs* alone. The frequency plot shows a skewed multimodal distribution where the majority of maps have moderate *EDs*≈*2* between them. This correspond to cases where both maps being compared show relatively simple, monotonic and smooth trait covariation (see example maps above in a normalized $T_1$-$T_2$ space). A second peak in frequency happens around *EDs*≈*6* account for the maps that, although still showing monotonic trait covariation (see above), exhibit flipped slopes (a situation that is statistically more common than no-correlated maps, see Fig. 2B). This third peak (*EDs*≈*8*.5) contains either highly non-linear maps or pairs of maps where one them shows no parametric sensitivity (a point in the $T_1$-$T_2$ space, see top right panel). This third peak roughly corresponds to the bulk of yellow off-diagonal points shown in (A). $n=10^6$. GRN + Multilinear model.

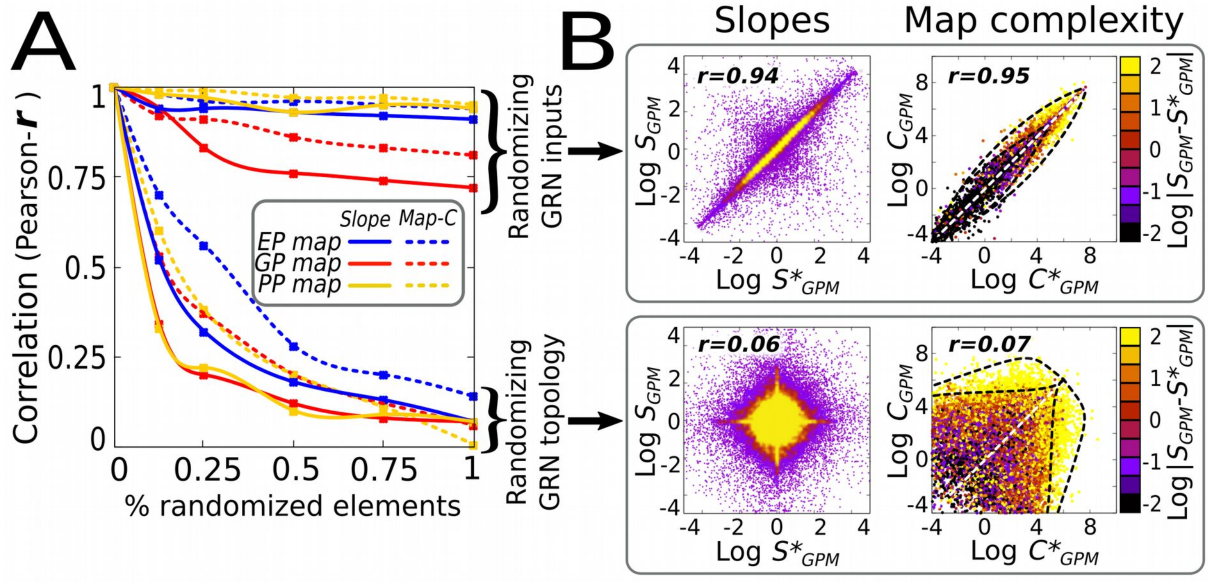


**Figure S4. GRN topology as the main contributor to phenotypic correlations.** The figure summarizes a stability test aimed to better understand the causal origins of the map-to-map correlations observed in Fig. 2 (specifically, if correlations arose from the developmental mechanism itself or from similarities on the input parameter values). In order to discriminate between these two causal factors, we gradually randomize both (with a probability $p=\{0.1,0.2,...1\}$, the GRN topology was changed as $M_{ij} \rightarrow |M_{ij}-1|$ and the GRN input parameters as $x \rightarrow x \sim U(0,1)$) while recording if the correlations between maps were still retained or not. Notice that in here correlations are established between maps arising from the unperturbed (elemental) GRN (slopes $S$ and map complexities $C$) and the perturbed, randomized ones ( $S^*$ and $C^*$). (A) For every GRN in our random ensemble, correlations are robust against input randomization but fall dramatically to non-significant values ($r<0.3$) when the GRN topology is allowed to change parametrically. The observed pattern is very similar for both slopes (solid lines) and map complexities (dashed lines). This insensitivity to the randomization in the input values (see SI1) suggests that they are not causally involved in the correlations observed in Fig. 2A. On the contrary, correlations rather show a strong dependence with the GRN topology, which embodies the dynamical developmental system that creates the phenotypes. (B) Each point plotted in (A) represents a correlation measured in a two-dimensional spaces for slopes $S$ or map complexities $C$ (As in Fig. 2A). We provide, as an illustrative example, two of these spaces corresponding to maps which arise from genetic perturbations (spaces for other types of perturbations are very similar, Fig. 2A). In the upper row, all the maps emerging from a set of genetic inputs $x=\{0,0.1,0.2,\ldots,1.0\}$ are compared with those emerging from maximally randomized genetic inputs $x \sim U(0,2)$. The limited scattering suggests that a given GRN produces almost identical GP maps irrespective of the inputs used. Below, using the same set of genetic inputs $x=\{0,0.1,0.2,\ldots,1.0\}$ and the same GRN ensemble, maps emerging from elemental GRNs and from perturbed GRNs whose topologies have been fully randomized ($M_{ij} \rightarrow |M_{ij}-1| \forall\ M_{ij}$) are compared, showing that, despite having identical inputs, they produce totally unrelated phenotypes. $r$ stands for Pearson's correlation. Yellow areas in $S_{GPM}$ *vs* $S^*_{GPM}$ spaces (left plots in B) contain 90% of GRNs. $n=10^6$. GRN + Multilinear model.

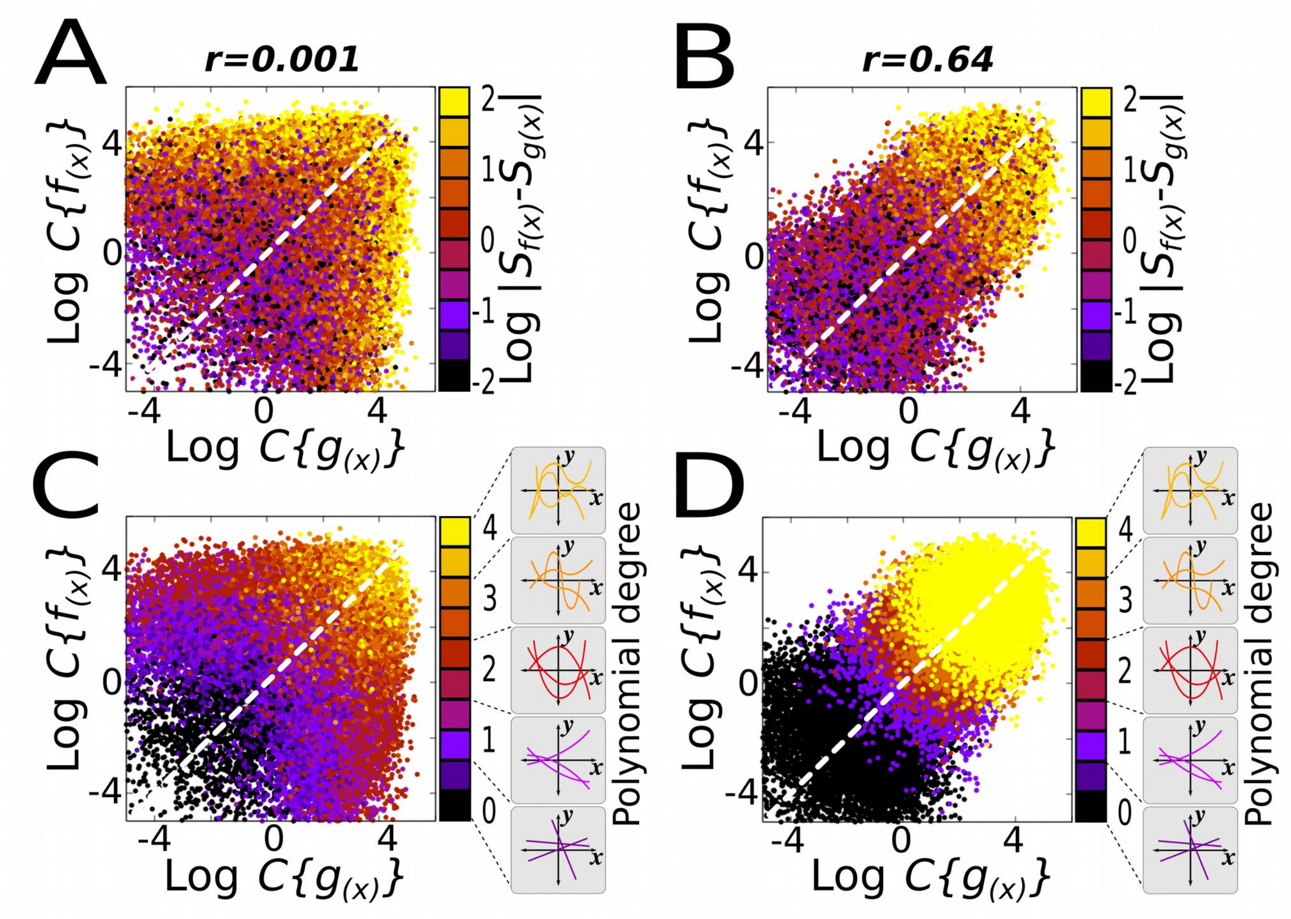


**Figure S5. Geometry alone does not explain the observed positive correlation between map complexities.** The aim of this experiment is to assess whether the observed correlation between map complexities (Fig. 2C) is a general feature mathematical functions or not. In order to do this, we analysed the relationship between function complexities (measured the same way as map complexity *C*, see SI1) in a large ensemble of random polynomial equations of known degree ($deg(f)\leq 4$), where the $x,y$ variables of $f(x)$ are considered to correspond to the $T_1$ and $T_2$ traits (see SI1). (A) When two functions $f(x)$ and $g(x)$ are randomly picked from the ensemble, no correlation between their map-complexities is observed. Colours express absolute differences between slopes (As in Fig. 2C). (B) A positive correlation between map complexities only appears when the functions to be compared are first sorted according to their complexity class, that is when $deg(g(x)):=deg(f(x))$. (C) The same as A, but colours now represent differences on the degree of the functions ($|deg(f(x))-deg(g(x))|$) rather than differences between slopes. (D) The same as A, but colours now denote the degree of the functions $deg(f(x))$ (since functions have been previously sorted, it is the same as $deg(g(x)$). Notice than in B and D, the correlation only appears as an inter-class effect simply because the function complexity correlates with the degree of that function ($deg(f(x)\alpha C\{f(x)\}$), but it is absent within the same class functions. This suggests that development creates correlations between the map complexities by restricting the number of developmental outcomes to well-defined and mechanism-specific possible general phenotypic distributions (GPDs, Fig. S1). $n=10^6$.

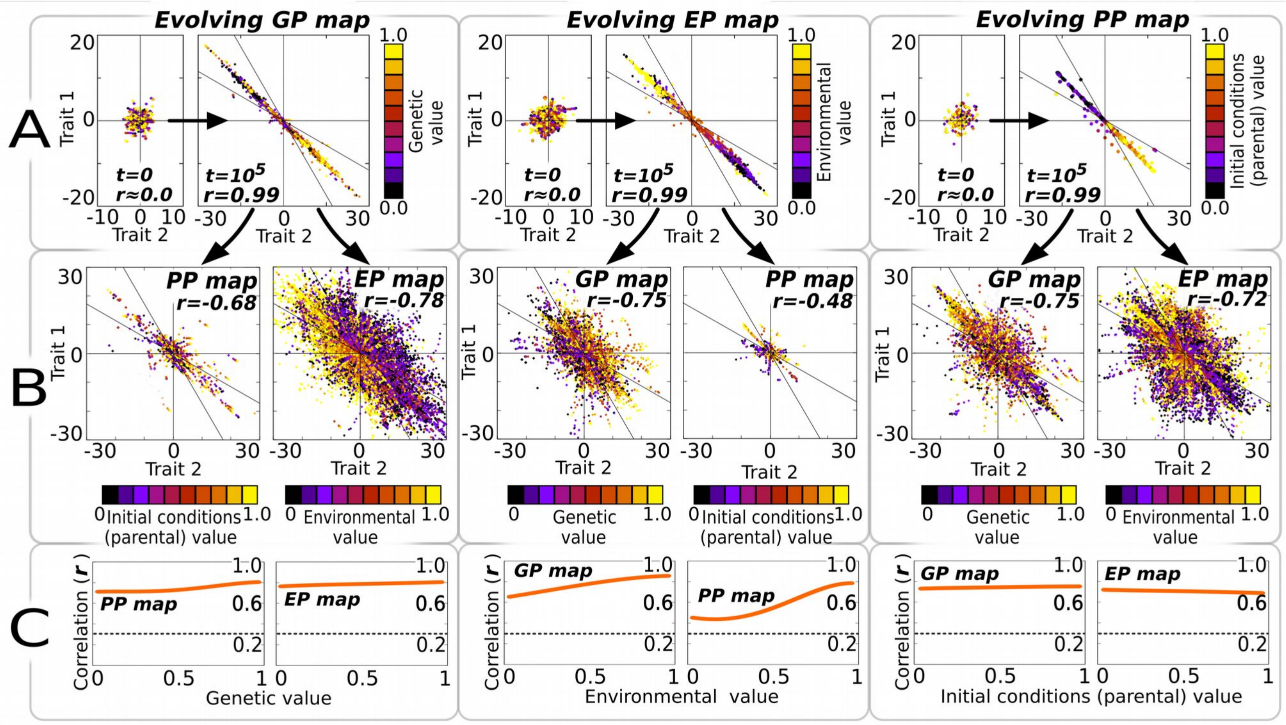


**Figure S6. Using negative slopes (anti-correlated traits) as a target does not change the general results of map evolution.** This figure reproduces the same results of Fig. 3, but in this case a target map with a slope *Sx=-1* (instead of *Sx=1*) has been used in the simulations. A comparison between both figures clearly reveals that results of the experiments involving adaptive evolution of maps are very similar when using different (linear) targets. As in Fig. 3, a population whose individuals initially exhibit no particular phenotypic distribution in *t=0* (A, small panels) is evolved to fit a target phenotypic distribution using as an input just one kind of phenotypic determinant. In each generation, one individual is exposed to 10 different values ($0<x<1$) of this phenotypic determinant, thus producing a set of ten potential phenotypes whose slope in a $T_1$-$T_2$ space is compared with the target to evaluate the individual's fitness $W_i$, see SI1). After $10^5$ generations in a mutation-selection-drift scenario (where other sources of phenotypic variation are frozen), population has a narrow phenotypic distribution in the evolved map (A, big panels). Then, we uncover the hidden variation in the other maps by i) freezing the variation in the parameter that generated the evolved map to an intermediate value and ii) introducing parametric variation ($0<x<1$) in the other phenotypic determinants (those that were kept fixed during the evolutionary trial). (B) Results reveal that evolving a single map adaptively creates significant side-effect phenotypic distributions in the other maps that are not the target of natural selection. (C) Correlations in the side-effect maps are significant irrespective of the value at which the parameter of the evolved map is frozen. $p=64$ individuals ; $n=30$ replicates. GRN + multilinear model.

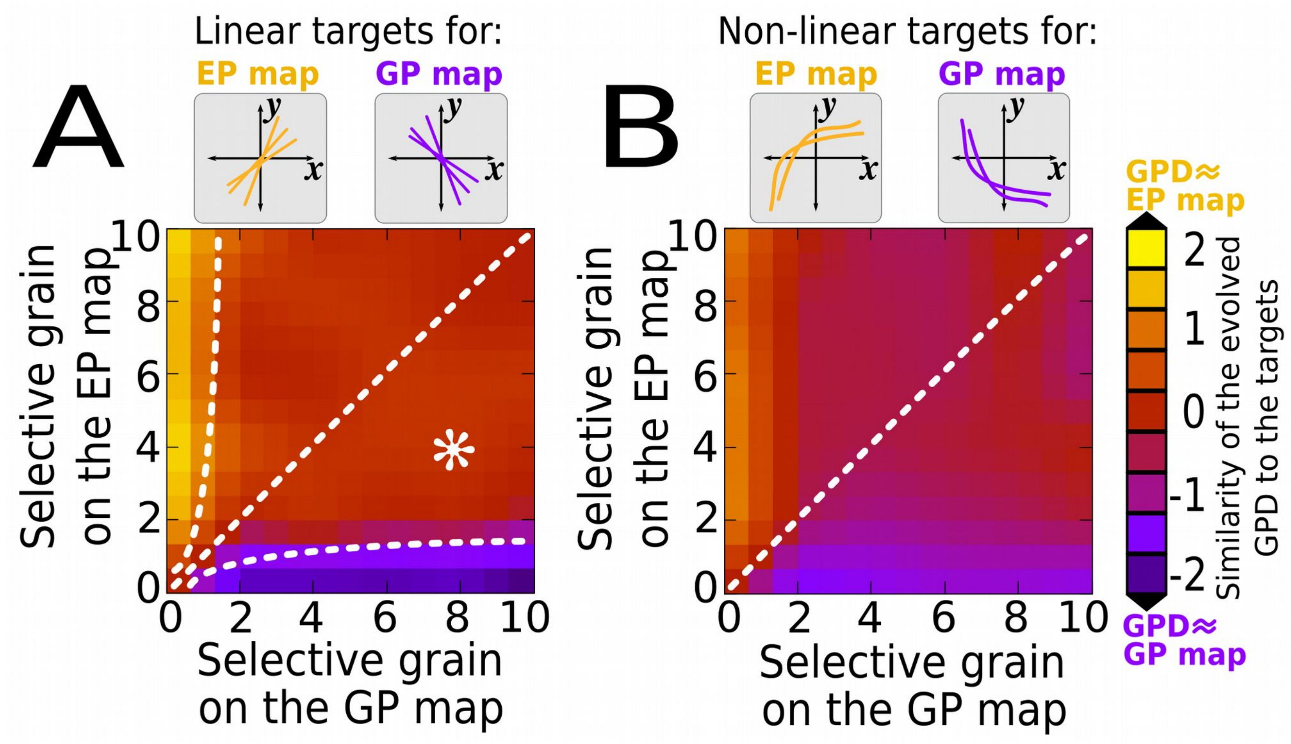


**Figure S7. The structure of the GPD is determined by the map under more fine-grained selection.** For this experiment, two different maps (EP map and GP map), each one having a different selective grain, are simultaneously evolved. (A) In order to discriminate between the effects of selection on the EP map and on the GP map, the slopes of the target maps are maximally different: $S^{T}_{EPM}=(S^{T}_{GPM})^{-1}$. Only the spatial component of fine-grainedness (how many "points" in the evolving maps are selectable, see SI1) is considered for this experiment. For instance, the white asterisk represents a population in which the fitness is calculated taking into account four points of the EPM and eight points of the GPM (in both cases, in each generation the these points are randomly selected from a set of 10). Colours represent the ED-based similarity of the general phenotypic distribution (GPD) to the targets after t=$10^4$ generations $ED(GPD,GPM_T)-ED(GPD,EPM_T)$. Basically, yellowish colour indicates that trait covariation found in the GPD closely resembles that of the $S^{T}_{EPM}$, and bluish colour that of the $S^{T}_{GPM}$. The resulting heatmap shows that the structure of the GPD is determined by the map under more fine-grained selection. The pattern is most clear when selection on one of the maps is very fine-grained (off-diagonal dashed lines) and in the other map is very coarse-grained. When the selective grain of the two maps is comparable, or when the selection on a map is not very fine-grained, intermediate maps not attributable to any target are found. (B) As in (A) but using non-linear targets (a random polynomial function with $deg(f)>2$). As Fig. 5 shows, the ability of the developmental systems to evolve non-linear maps is severely limited, so here most of the evolved maps are different from both targets (if the target map is too non-linear, nor the genotype nor the environment can be the leaders of adaptive change). $n=30$ replicates, $p=64$ individuals, GRN + multilinear model.

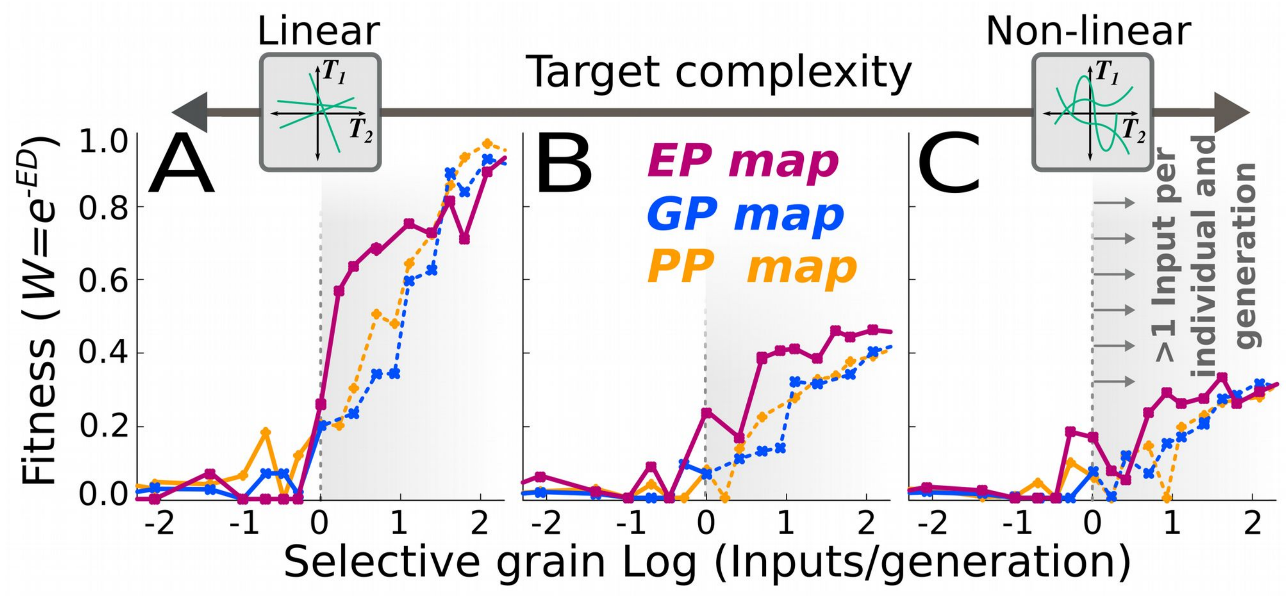


**Fig. S8. Simpler maps evolve more easily than complex ones.** As in Fig. 5, this figure shows the efficiency of natural selection in evolving certain maps (fitness achieved, vertical axis) under different regimes of selective grain (the average number of parameter-phenotype points that can be "seen" by natural selection in each generation, horizontal axis). In this experiment, however, we use also different non-linear functions (random polynomials) as targets maps. (A) shows the same result shown in Fig. 5: Linear maps can be easily evolved given certain degree of selective grain. (B-C) generalise this result to non-linear target functions: complex maps are hard to evolve even under fine-grained selection regimes. This suggests that the adaptive evolution of very complex maps would require biologically unrealistic levels of selective grain (although they could be attained by means of non-adaptive mechanisms, see main text). For a given target complexity, specially for simpler ones, maximal efficiency is generally achieved when single individuals can experience more than one input per generation ($Log_{(Input/Generation)}>0$, shadowy areas). Biologically, these high levels of selection fine-grainedness can only be achieved by the Environment-to-Phenotype (EP) map (see main text and Fig. 5). For the sake of clarity, only averages over the $n=30$ replicates are shown (points), and Log-scale is used in the horizontal axis. Euclidean-distance (*ED*)-based fitness. $p=64$ individuals; $t=10^4$ generations, GRN + Multilinear model. For each replicate, the target map is a polynomial function of known degree (complexity) and arbitrary non-zero coefficients.
